# Ncf1/Igfbp2 Signaling Regulates Airway Submucosal Gland Progenitor Fitness and Ionocyte Specification

**DOI:** 10.64898/2026.10.13.758956

**Authors:** Javahir Hajibabazade, Kyhler D. Campbell, Oluwadarasimi Atanda, Grace Fritz, Charles A. Yeaman, Weam S. Shahin, John F. Engelhardt

## Abstract

**Background:** Airway submucosal gland progenitors (SMGPs) serve as reserve stem cells for epithelial regeneration, yet mechanisms governing their long-term maintenance remain poorly understood. Here, we identify Ncf1/Igfbp2 signaling as a regulator of mouse SMGP fitness and epithelial lineage specification.

**Methods:** Mouse SMGPs with genetic disruption of *Ncf1* or *Igfbp2* were serially passaged to assess progenitor proliferation, organoid-forming capacity, epithelial marker expression, DNA damage, DNA repair responses, and telomere length. Differentiated epithelia were evaluated for Cftr-mediated anion transport and pulmonary ionocyte specification in vitro and in vivo.

**Results:** Genetic disruption of *Ncf1* or *Igfbp2* preserved SMGP identity, enhanced proliferative and organoid-forming capacity, and reduced DNA damage accumulation through activation of DNA repair programs. Epithelia derived from *Ncf1*^−/−^ and *Igfbp2*^−/−^ SMGPs maintained higher Cftr-mediated anion transport at late passage and exhibited enhanced pulmonary ionocyte specification in vitro and in vivo. These effects occurred independently of telomere shortening, implicating altered stress responses and DNA damage handling rather than replicative limits.

**Conclusions:** Together, our findings identify Ncf1/Igfbp2 signaling as a regulator of airway stem cell fitness and epithelial maturation, with implications for regenerative therapies in lung diseases such as cystic fibrosis.

## Introduction

Airway stem cells are emerging as promising therapeutic targets in pulmonary regenerative medicine for diseases including pulmonary fibrosis, chronic obstructive pulmonary disease, acute respiratory distress syndrome, and genetic disorders such as primary ciliary dyskinesia and cystic fibrosis (CF). This therapeutic potential stems from their capacity for self-renewal, lineage-specific differentiation, restoration of epithelial integrity, and tissue repair. Combined with gene editing, airway stem cells may also provide durable treatments for inherited disorders such as CF. Beyond cell-based therapies, airway stem cells are widely used for disease modeling, organoid studies, drug screening, and tissue engineering [1-6]. Despite their promise, major challenges remain, including stem cell exhaustion [7] and progressive loss of multipotency [8] during long-term expansion.

Reactive oxygen species (ROS) regulate stem cell self-renewal, proliferation, differentiation, senescence, and apoptosis [9-12]. Cellular ROS arise from both exogenous and endogenous sources, including mitochondria and NADPH oxidases [13]. Among the seven NADPH oxidase family members [14], Nox2 is particularly notable for generating superoxide through electron transfer from NADPH to molecular oxygen [15]. Nox2 activation requires assembly of cytosolic subunits Ncf1 (p47phox), p40phox, p67phox, and Rac1 with membrane-bound p22phox and gp91phox. Genetic disruption of *Ncf1* in mice revealed that Nox2-dependent ROS increases Igfbp2 bioavailability, thereby regulating neural stem cell self-renewal and lineage commitment through DNA repair pathways [16]. Phagosomal Nox2-derived superoxide also contributes to tissue inflammation [15], including CF lung disease [17, 18].

ROS signaling has also been implicated in airway epithelial homeostasis. Dynamic intracellular ROS flux regulates airway basal stem cell self-renewal and proliferation through Nrf2-dependent activation of Notch signaling. Moderate ROS levels support basal stem cell homeostasis, whereas restoration of redox balance through antioxidant programs is required to maintain epithelial stability [9-12]. Although these findings establish ROS as an important regulator of airway stem cell behavior, the contribution of specific Nox2-associated redox pathways to long-term progenitor maintenance and lineage specification remains poorly understood.

Among airway stem cell populations, submucosal gland progenitors (SMGPs) are of particular interest because they function as reserve stem cells capable of regenerating both submucosal glands and surface airway epithelium following severe injury [19, 20]. In this study, we investigated the role of Ncf1/Igfbp2 signaling in regulating SMGP stemness and multipotency using genetic disruption of both genes in mice coupled with extensive serial passaging. Disruption of this signaling axis preserved progenitor characteristics during long-term expansion, reduced DNA damage accumulation, and increased expression of DNA repair genes. Furthermore, interruption of Ncf1/Igfbp2 signaling promoted pulmonary ionocyte enrichment and enhanced Cftr-mediated ion transport, identifying a redox-associated stress pathway that regulates airway progenitor state and epithelial function.

## Results

### Loss of Ncf1/Igfbp2 signaling reduces DNA damage and upregulates DNA repair programs in high-passage SMGPs

To determine whether Ncf1/Igfbp2 signaling influences SMG progenitor behavior, we evaluated the expression of the airway basal cell markers Keratin 5 (Krt5) and the DNA double-strand break marker γ-H2ax following depletion of *Ncf1* or *Igfbp2*. Krt5 was selected based on evidence that SMGPs adopt a basal cell-like phenotype in culture [19, 20]. γ-H2ax expression was examined because Ncf1/Igfbp2 signaling has been previously implicated in regulating neural stem cell fate through modulation of the DNA damage response [16]. To model replicative senescence, we employed serial passaging in an in vitro system that enables investigation of cellular and molecular changes associated with stem cell aging, including exhaustion and loss of multipotency.

SMGPs were isolated from 8-week-old wild-type (WT) C57BL/6J, *Igfbp2*^*−/−*^, and *Ncf1*^*−/−*^ mice, and passages 4 (P4) and 19 (P19) were harvested for assessment of Krt5 and γ-H2ax expression by western blot (WB). From P4 to P19, WT SMGPs exhibited a marked downregulation of Krt5, whereas both *Igfbp2*^*− /−*^ and *Ncf1*^*−/−*^ SMGPs retained high levels of Krt5 expression at late passage (P19) **(Fig. 1A, B, D, E)**. Conversely, γ-H2ax expression was significantly increased in WT SMGPs over this interval but remained relatively low in *Igfbp2*^*−/−*^ and *Ncf1*^*−/−*^ SMGPs **(Fig. 1A, C, D, F)**, suggesting less accumulation of DNA damage in *Igfbp2*^*−/−*^ and *Ncf1*^*−/−*^ SMGPs compared to WT counterparts. To further determine whether these differences were associated with altered DNA repair capacity, we assessed by quantitative real-time PCR (qPCR) the expression of key Fanconi anemia (FA) pathway genes, including *Fanca, Fancd2*, and *Rad51*. These genes were selected based on their central roles in homologous recombination [21] and interstrand cross-link repair [22], processes implicated in the DNA damage response pathways regulated by Ncf1/Igfbp2 signaling in neural stem cells [16]. While no significant differences in *Rad51, Fanca*, or *Fancd2* expression were observed at P4 **(Fig. 1G)**, both *Igfbp2*^*−/−*^ and *Ncf1*^*−/−*^ SMGPs exhibited significantly higher expression of these DNA repair genes at P19 relative to WT controls **(Fig. 1H)**. Together, these findings suggest that Ncf1/Igfbp2 signaling modulates DNA damage responses that influence the maintenance and functional capacity of SMG basal-like progenitors during in vitro expansion.

**Figure 1.**
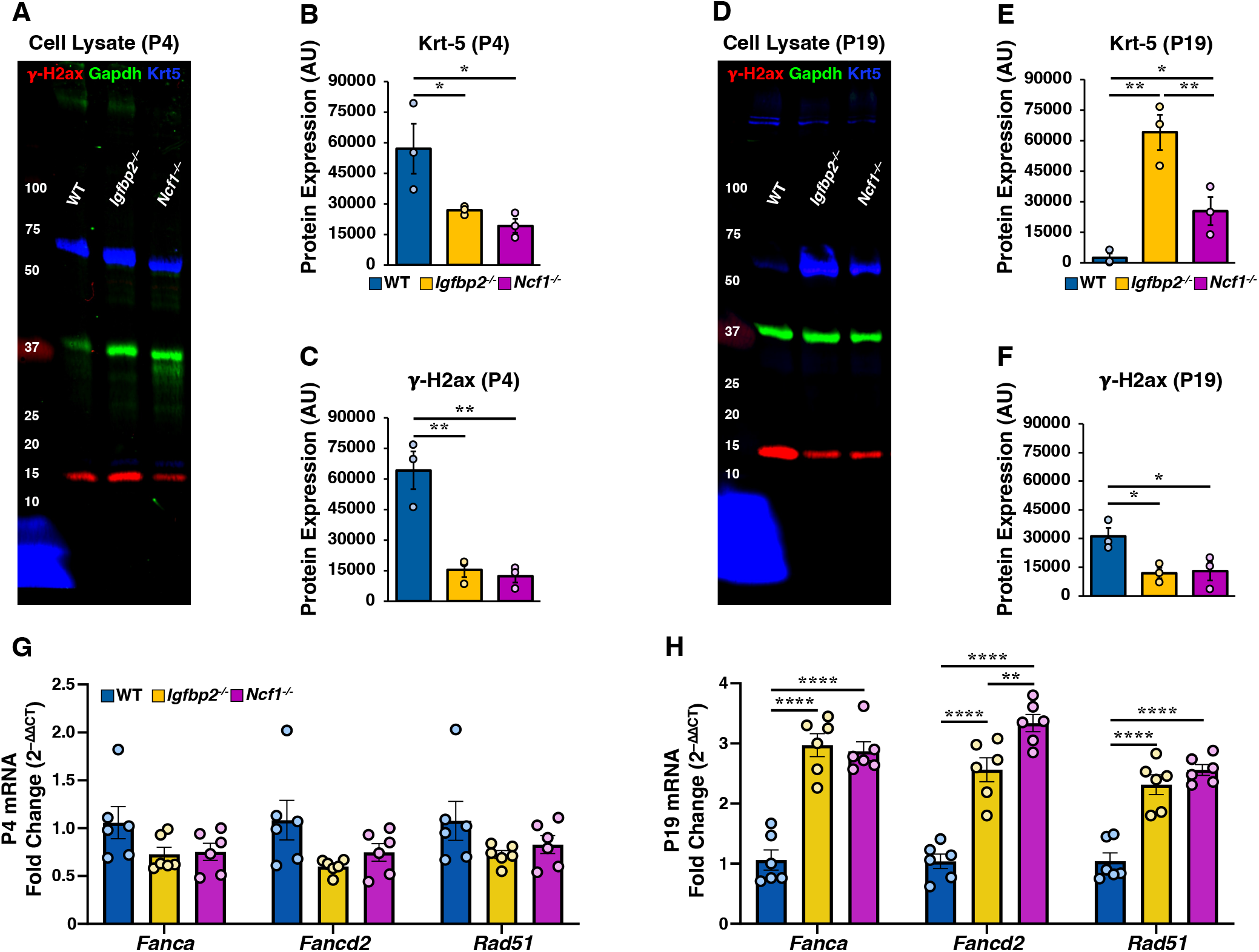
Reduced DNA damage and increased DNA repair gene expression in Ncf1/Igfbp2-deficient SMG progenitors at high passage. **(A, D)** Representative western blots of Krt5 and γ-H2ax in WT, *Igfbp2*^*−/−*^, and *Ncf1*^−/−^ SMG progenitors at passage 4 (A) and 19 (D). **(B, E)** Quantification of Krt5 expression normalized to Gapdh. **(C, F)** Quantification of γ-H2ax expression normalized to Gapdh (n = 3 cultures per cell type). **(G, H)** qPCR analysis of *Fanca, Fancd2*, and *Rad51* in proliferating SMG progenitors at P4 (G) and P19 (H) (n = 6 cultures per genotype, with 2 cultures from each of 3 donor groups; each donor group comprised SMGPs from 3 mice). Gene expression was normalized to *Gapdh*, and relative expression was calculated using the 2^−ΔΔCT^ method, with ΔΔCT values determined relative to the mean ΔCT of WT controls. Data are shown as mean ± SEM. Statistical significance: *P < 0.05, **P < 0.01, ***P < 0.001, ****P < 0.0001; one-way ANOVA with Benjamini−Hochberg FDR post hoc test.

### Genetic disruption of Ncf1/Igfbp2 signaling is implicated in SMGP proliferation and organoid-forming capacity throughout serial passaging in culture

To determine whether Ncf1/Igfbp2 signaling regulates the proliferative capacity of SMGPs during long-term expansion, EdU incorporation assays were performed using a 3-hour pulse followed by either immediate (0h) or 24-hour chase at early (P4) and late (P19) passages **(Suppl. Fig. S1A, B)**. At P4, robust EdU incorporation was observed across all genotypes, and quantification revealed no significant differences in the percentage of EdU-positive cells (EdU^+^) at either time point **(Suppl. Fig. S1C)**. The increase in EdU^+^ cells from 0h to 24h was similarly maintained across WT, *Igfbp2*^*−/−*^ and *Ncf1*^*−/−*^ cultures, indicating comparable proliferative capacity at early passage. In contrast, marked differences in EdU incorporation were detected at P19 **(Suppl. Fig. S1D)**. Both *Igfbp2*^*−/−*^ and *Ncf1*^*−/−*^ SMGPs exhibited significantly higher percentages of EdU^+^ cells compared to WT counterparts at both 0 h and 24 h, with *Igfbp2*^*−/−*^ SMGPs displaying the highest proliferative index. These findings suggest that disruption of Ncf1 or Igfbp2 preserves proliferative capacity during extended in vitro expansion. To further evaluate progenitor activity and self-renewal, organoid-forming efficiency (OFE) was assessed at P4 and P19 **(Suppl. Fig. S1E)**. No significant differences in OFE were observed among genotypes at P4. However, P19 *Igfbp2*^*−/−*^ SMGPs exhibited the highest OFE relative to WT and *Ncf1*^*−/−*^ cultures, as well as to their corresponding P4 controls. P19 *Ncf1*^*−/−*^ SMGPs also exhibited significantly higher OFE compared to WT, although not as great as *Igfbp2*^*−/−*^ cultures. The relative OFE across genotypes correlated with Krt5 protein expression **(Fig. 1E)**, in which P19 *Igfbp2*^*−/−*^ SMGPs exhibited the highest Krt5 expression, *Ncf1*^*−/−*^ cultures displayed intermediate expression, and WT SMGPs showed the lowest levels. Together, these findings suggest that loss of *Ncf1* or *Igfbp2* preserves basal-like progenitor features and supports long-term self-renewal during in vitro expansion.

### Ncf1/Igfbp2 signaling influences long-term expansion, basal cell marker expression, and telomere dynamics in airway SMGPs

To determine whether Ncf1/Igfbp2 signaling regulates long-term SMGP maintenance, cell yield was quantified during serial passaging from P4 to P19 **(Suppl. Fig. S2A)**. No differences in cell yield were observed during early passages. However, beginning at P14, WT SMGPs exhibited reduced expansion relative to *Igfbp2*^*−/−*^ cultures. By P17 and P19, WT SMGPs displayed significantly lower cell yield relative to both *Igfbp2*^*−/−*^ and *Ncf1*^*−/−*^ cultures. At P19, *Ncf1*^*−/−*^ SMGPs exhibited intermediate expansion between *Igfbp2*^−/−^ and WT cultures. These findings suggest that loss of Ncf1 or Igfbp2 preserves long-term expansion during extended culture. *Krt5* expression was also comparable across genotypes at P4 but was significantly lower in WT cultures relative to both knockout groups at P19 **(Suppl. Fig. S2B)**, consistent with the Krt5 protein expression patterns **(Fig. 1D)**. Because Krt14 expression is associated with airway injury and stress-responsive remodeling across multiple disease models and injury conditions [19, 23, 24], we examined *Krt14* expression and found that it did not differ across genotypes at P4 but was significantly elevated in WT cultures at P19 relative to *Igfbp2*^*−/−*^ and *Ncf1*^*−/−*^ SMGPs **(Suppl. Fig. S2C)**.

*Pcna* expression was next evaluated by qPCR. Although *Pcna* levels were comparable across genotypes at P4, expression was significantly increased in WT cultures at P19 relative to both knockout groups **(Suppl. Fig. S2D)**. Pcna functions as a DNA polymerase processivity factor during DNA replication and also participates in DNA damage tolerance pathways. Under conditions of replication stress, PCNA undergoes post-translational modifications that facilitate lesion bypass by recruiting translesion synthesis polymerases and coordinating DNA damage tolerance responses [25-27]. The elevated Pcna expression in late-passage WT cultures, together with increased γ-H2AX and reduced expression of Fanconi anemia pathway DNA repair genes, may reflect increased replication-associated stress and activation of DNA damage tolerance pathways. Collectively, these findings further support the notion that WT SMGPs experience increased cellular stress during long-term expansion, whereas *Igfbp2*^− /−^ and *Ncf1*^−/−^ cultures maintain progenitor integrity through reduced DNA damage accumulation and increased expression of DNA repair genes.

To determine whether differences in long-term expansion were associated with altered telomere dynamics, absolute telomere length per chromosomal end was quantified **(Suppl. Fig. S2E)**. WT SMGPs exhibited significantly longer telomeres than *Igfbp2*^−/−^ and *Ncf1*^−/−^ cultures across all passages examined (P4, P10, P15, and P19). However, normalization to baseline (P4) revealed no differences in relative telomere shortening across genotypes **(Suppl. Fig. S2F, G)**, indicating comparable rates of telomere attrition during serial passaging. To further evaluate telomere regulatory pathways, expression of *Terf2*, a key component of the shelterin complex that mediates telomere capping and protection from DNA damage signaling, and *Tert*, the catalytic subunit of telomerase, was assessed. *Terf2* expression was significantly higher in WT cultures at P4 relative to both knockout groups but did not differ at P19 **(Suppl. Fig. S2H)**. *Tert* expression remained comparable across genotypes at both passages **(Suppl. Fig. S2I)**. Together, these findings suggest that enhanced expansion of *Igfbp2*^−/−^ and *Ncf1*^−/−^ SMGPs is not associated with differences in the rate of telomere attrition but instead reflects differences in cellular stress responses and DNA damage handling during extended culture.

### Disruption of Ncf1/Igfbp2 signaling is associated with reduced oxidative stress in long-term-cultured SMG progenitors

Given the established role of Ncf1 in regulating NADPH oxidase, we next examined whether differences in oxidative stress accompanied the altered progenitor fitness observed during long-term SMGP expansion. Ncf1-dependent ROS production has been implicated in redox modification of Igfbp2 and in the regulation of DNA repair pathways and neural stem cell fate [16]. In airway basal stem cells, dynamic increases in intracellular ROS from low to moderate levels have also been shown to support self-renewal and proliferation [9]. We therefore assessed intracellular oxidative stress in live WT, *Igfbp2*^−/−^, and *Ncf1*^−/−^ SMGPs using CellROX Deep Red dye at P4 and P19 (**Fig. 2A, B**). CellROX Deep Red was chosen because it is highly selective for superoxide and hydroxyl radical and would partially inform NADPH oxidase involvement. At P4, CellROX fluorescence intensity was comparable across genotypes (**Fig. 2C**). In contrast, at P19, both *Igfbp2*^−/−^ and *Ncf1*^−/−^ cultures maintained significantly lower CellROX fluorescence than P19 WT cultures (**Fig. 2D**). Thus, genotype-dependent differences in oxidative stress emerged at late passage and coincided with the reduced DNA damage accumulation and enhanced progenitor fitness observed in *Igfbp2*^−/−^ and *Ncf1*^−/−^ SMGPs.

**Figure 2.**
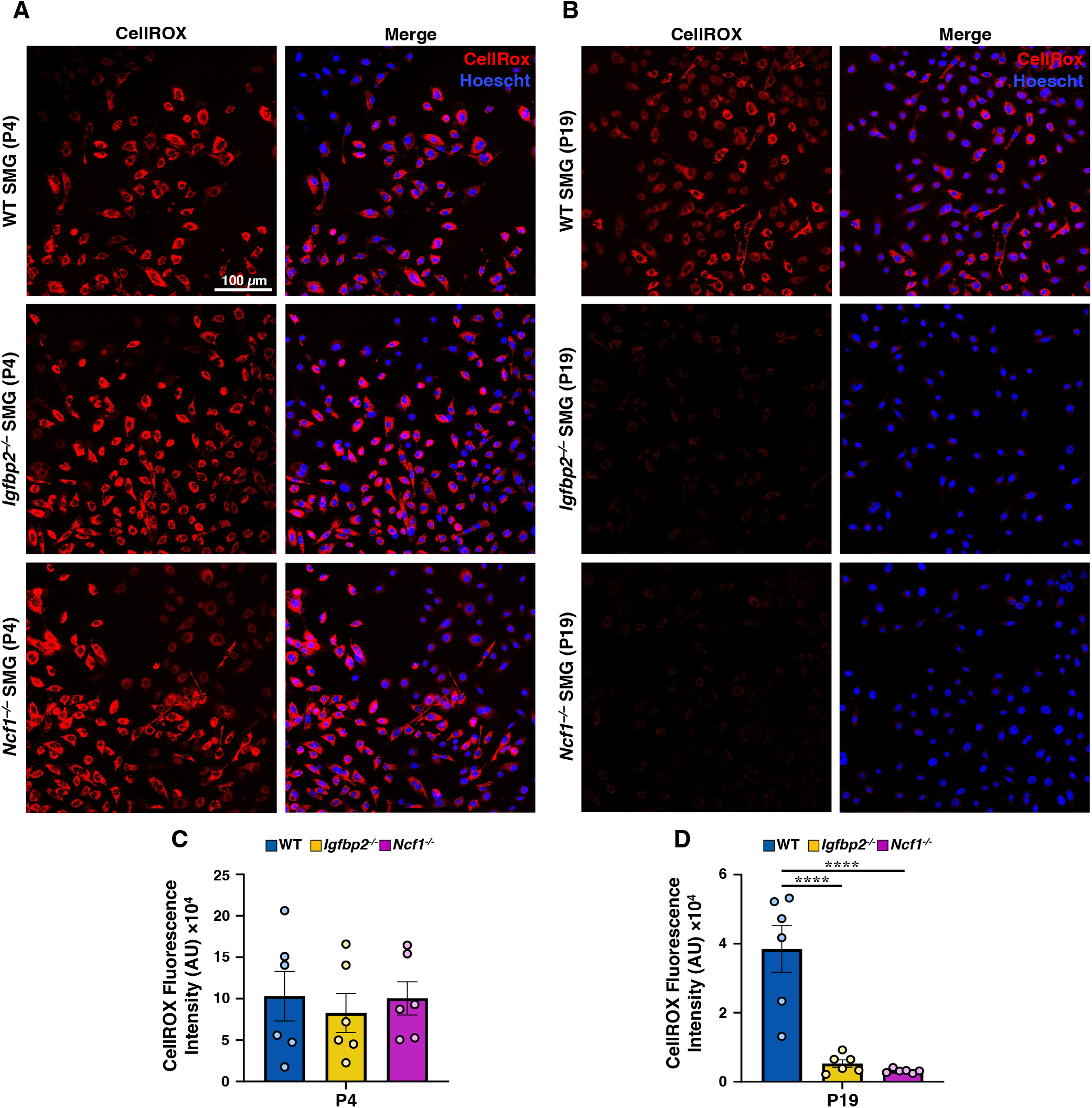
Long-term SMGP expansion is associated with genotype-dependent differences in oxidative stress. **(A, B)** Representative images of CellROX Deep Red fluorescence (red, left) and merged CellROX (red) and Hoechst (blue) fluorescence (right) in live WT, *Igfbp2*^−/−^, and *Ncf1*^−/−^ SMG progenitors at P4 (A) and P19 (B) following a 30-min incubation with CellROX and a subsequent 10-min incubation with Hoechst. **(C, D)** Quantification of CellROX fluorescence intensity at P4 (C) and P19 (D) (n = 6 cultures per genotype, with 2 cultures from each of 3 donor groups; each donor group comprised SMGPs from 3 mice). Data are presented as mean ± SEM. Statistical significance: *P < 0.05, **P < 0.01, ***P < 0.001; one-way ANOVA with Benjamini−Hochberg FDR post hoc test.

### Loss of Ncf1/Igfbp2 signaling preserves functional anion transport in serially passaged SMGPs

Previous studies have demonstrated that serial passaging of airway basal cell progenitors from multiple species leads to a progressive loss of CFTR-mediated anion transport following polarization at an air-liquid interface (ALI) [28]. This reduction has been attributed to the loss of multipotency and reduced capacity to differentiate into CFTR-expressing epithelial cell types following extended serial passaging [29-33]. We therefore hypothesized that diminished pulmonary ionocyte specification contributes to this functional decline and that disruption of Ncf1/Igfbp2 signaling would preserve ionocyte differentiation and Cftr function in late-passage cultures. To test this hypothesis, we established the polarized ALI cultures of WT, *Igfbp2*^*−/−*^, and *Ncf1*^*−/−*^ SMGPs at P4 and P19. Fully differentiated ALI cultures were analyzed by Ussing chamber electrophysiology to measure short circuit current (*Isc*). Under symmetrical chloride conditions, cAMP-inducible and GlyH101/CFTR_Inh_172-inhibitable Cftr-mediated secretory chloride currents were significantly higher in *Ncf1*^*−/−*^ and *Igfbp2*^*−/−*^ SMGP-derived ALI cultures relative to WT controls at P4 **(Fig. 3A, C)**. This pattern was maintained at P19, where *Ncf1*^*−/−*^ and *Igfbp2*^*−/−*^ ALI cultures continued to exhibit significantly higher Cftr-mediated chloride currents than WT controls **(Fig. 3B, D)**. These results demonstrate that loss of Ncf1/Igfbp2 signaling preserves the ability of serially passaged SMGPs to generate polarized epithelia with robust Cftr-mediated anion transport.

**Figure 3.**
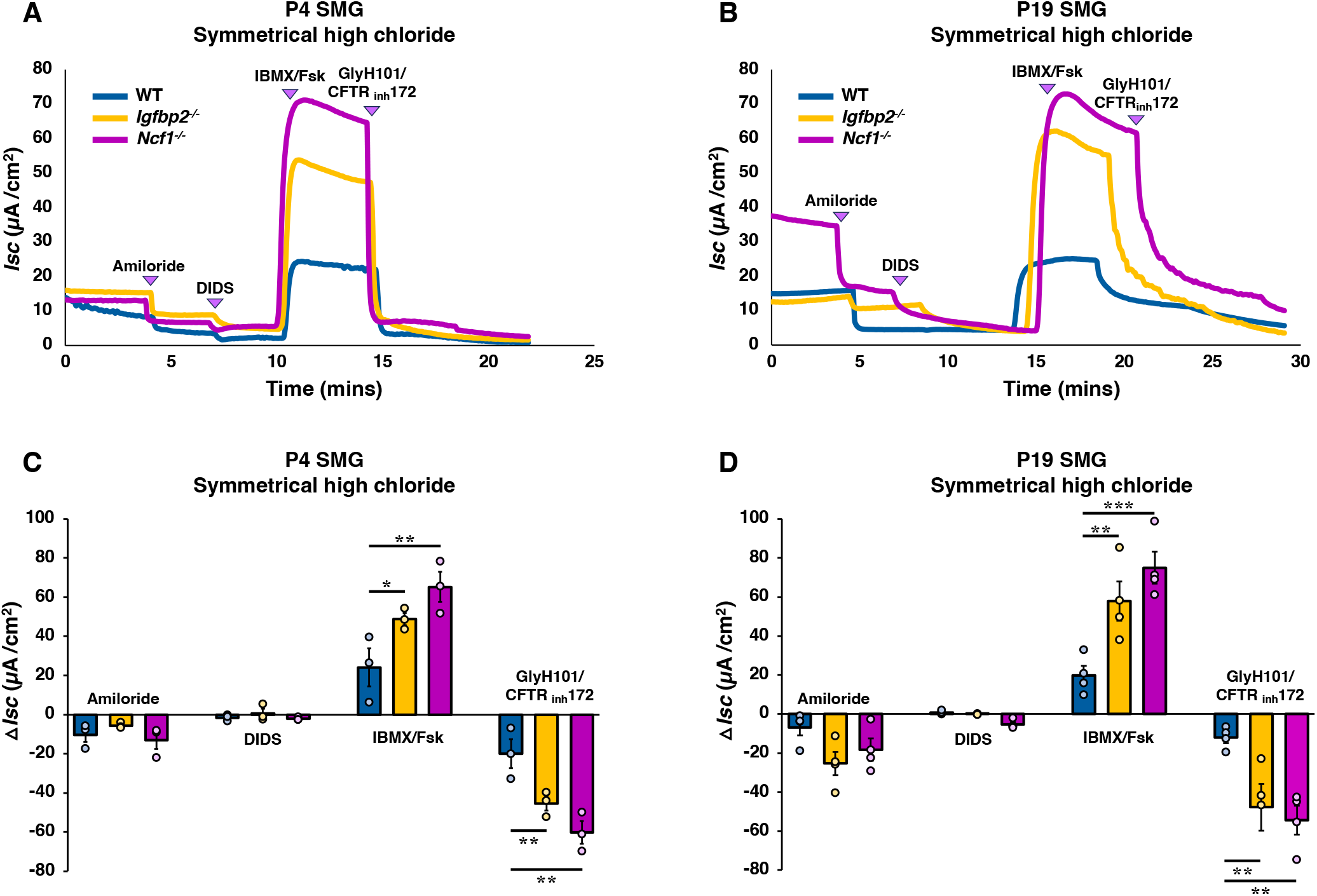
Polarized ALI cultures derived from SMG progenitors with Ncf1/Igfbp2 disruption maintain the elevated transepithelial anion transport through extensive passaging. **(A, B)** Representative short-circuit current (*Isc*) traces recorded in Ussing chambers following sequential addition of amiloride, DIDS, IBMX/forskolin, and GlyH101/CFTR_Inh_172 in WT, *Igfbp2*^−/−^, and *Ncf1*^−/−^ SMG-derived ALI cultures at passage 4 (P4) (A) and passage 19 (P19) (B). **(C, D)** Quantification of changes in chloride short-circuit current (Δ*Isc*) in response to amiloride, DIDS, IBMX/forskolin, and GlyH101/CFTR_Inh_172 in WT, *Igfbp2*^−/−^, and *Ncf1*^−/−^ SMG-derived ALI cultures at P4 (C; n = 3 cultures per condition) and P19 (D; n = 4 cultures per condition). Data are shown as mean ± SEM. Statistical significance: *P < 0.05, **P < 0.01, ***P < 0.001, ****P < 0.0001; one-way ANOVA with Benjamini− Hochberg FDR post hoc test.

### Loss of Ncf1/Igfbp2 signaling promotes pulmonary ionocyte specification

In the airway epithelium, CFTR is expressed predominantly in secretory cells and pulmonary ionocytes [34-37]. The discovery of CFTR-rich pulmonary ionocytes prompted considerable interest in their contribution to airway ion and fluid transport [5, 34, 37, 38]. Because disruption of Ncf1/Igfbp2 signaling enhanced Cftr-mediated currents in differentiated ALI cultures at both early and late passages, we next investigated whether this phenotype was associated with altered pulmonary ionocyte or secretory cell specification. RNA was isolated from WT, *Igfbp2*^*−/−*^, and *Ncf1*^*−/−*^ SMGP-derived ALI cultures at P4 and P19 for qPCR. *Ncf1*^*−/−*^ and *Igfbp2*^*−/−*^ ALI cultures exhibited significantly higher expression of ionocyte marker transcripts (*Foxi1, Bsnd* and *Cftr*) relative to WT cultures at both P4 and P19 **(Fig. 4A−C, F−H)**. The expression levels of ciliated and secretory cell markers, *FoxJ1* and *Spdef*, respectively, did not differ significantly across genotypes at P4 **(Fig. 4D, E)**. In contrast, P19 ALI cultures derived from *Igfbp2*^*−/−*^ and *Ncf1*^*−/−*^ SMGPs exhibited significantly increased expression of *FoxJ1* and *Spdef*, respectively, relative to WT cultures (**Fig. 4I, J)**. These findings suggest that enhanced Cftr-mediated chloride transport following disruption of Ncf1/Igfbp2 signaling is associated with increased pulmonary ionocyte specification.

**Figure 4.**
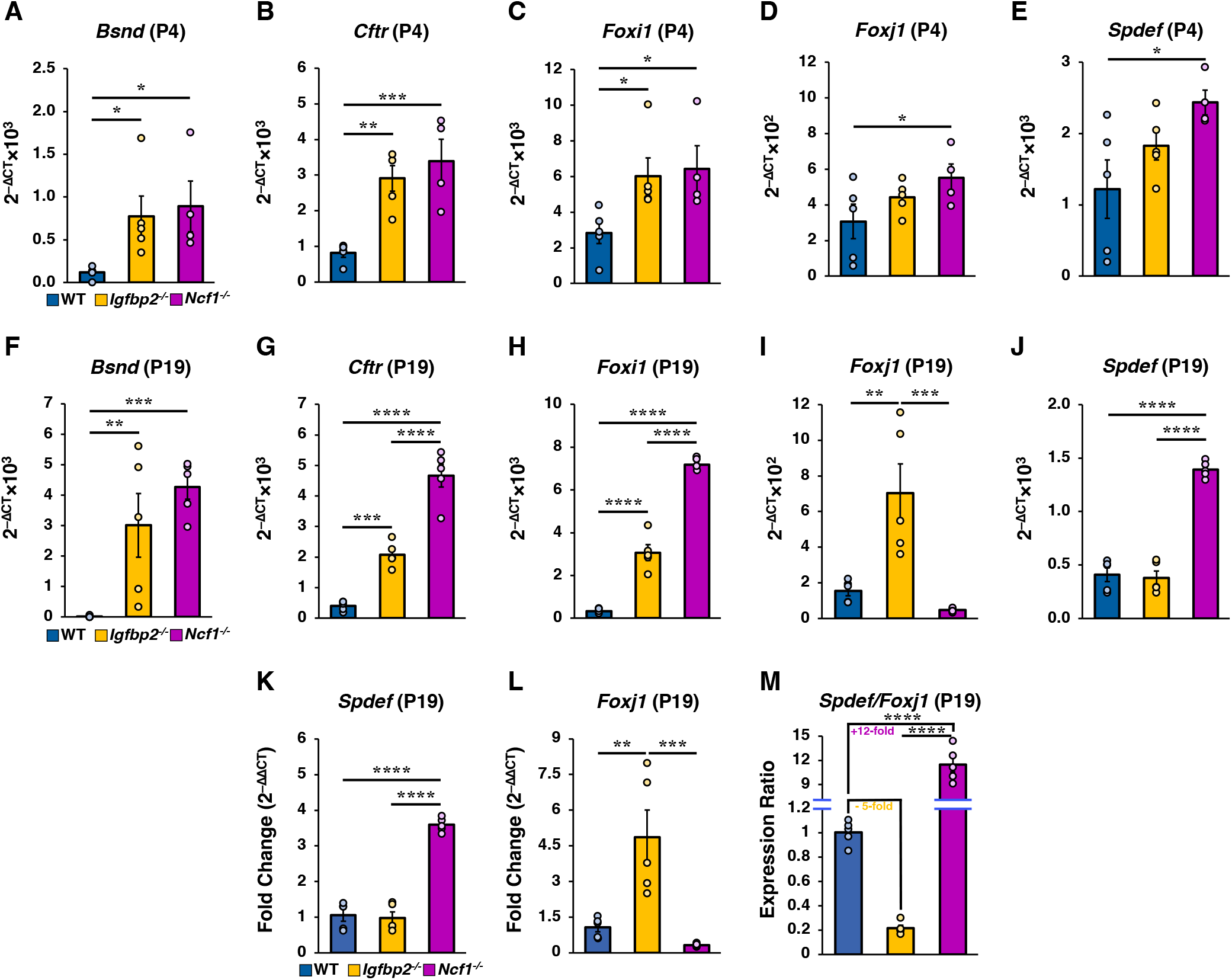
Disruption of Ncf1/Igfbp2 signaling promotes and maintains the expression of ionocyte markers. **(A−J)** Total RNA was isolated from polarized WT, *Igfbp2*^*−/−*^, and *Ncf1*^*−/−*^ SMG-derived ALI cultures at passage 4 (P4) and passage 19 (P19), reverse transcribed into cDNA, and analyzed by qPCR. Expression levels of *Bsnd* **(A, F)**, *Cftr* **(B, G)**, *Foxi1* **(C, H)**, *Foxj1* **(D, I)**, and *Spdef* **(E, J)** were measured in WT, *Igfbp2*^*−/−*^, and *Ncf1*^*−/−*^ cultures at P4 (A−E) and P19 (F−J) (n = 5 independent cultures per group unless otherwise indicated; *Ncf1*^*−/−*^ cultures at P4 were analyzed with n = 4 cultures). Gene expression was calculated using the 2^−ΔCT^ method and presented as 2^−ΔCT^ × 10^2^ or 2^−ΔCT^ × 10^3^ following normalization to housekeeping gene expression. **(K−L)** Fold-change expression of *Spdef* (K) and *Foxj1* (L) in P19 cultures was calculated using the 2^−ΔΔCT^ method relative to WT controls. **(M)** *Spdef*/*Foxj1* expression ratio in P19 cultures, calculated as 2^−(ΔΔCT*Spdef* − ΔΔCT*Foxj1*)^. Data are shown as mean ± SEM. Statistical significance: *P < 0.05, **P < 0.01, ***P < 0.001, ****P < 0.0001; one-way ANOVA with Fisher’s LSD post hoc test.

To further examine lineage-associated transcriptional differences at late passage, ΔΔCt analysis was performed for *Spdef* and *Foxj1* relative to WT controls **(Fig. 4K, L)**. This analysis revealed distinct lineage-associated transcriptional profiles between knockout genotypes, with *Ncf1*^−/−^ cultures exhibiting higher *Spdef* expression and *Igfbp2*^−/−^ cultures exhibiting higher *Foxj1* expression. To quantify these relative shifts, we calculated the *Spdef*/*Foxj1* expression ratio as a surrogate measure of secretory versus ciliated lineage bias. Compared with WT cultures, this ratio was increased ∼15-fold in *Ncf1*^−/−^ epithelia and decreased ∼5-fold in *Igfbp2*^−/−^ epithelia **(Fig. 4M)**. Together, these findings suggest that Ncf1/Igfbp2 signaling influences epithelial lineage allocation beyond pulmonary ionocytes and may differentially regulate secretory and ciliated cell differentiation programs during long-term expansion.

Because pulmonary ionocytes and secretory cells can both express Cftr and arise from interconnected epithelial differentiation programs, we next examined how disruption of Ncf1/Igfbp2 signaling affected cell type-specific Cftr expression. To validate our transcriptional findings at the protein level, whole mount immunofluorescence (IF) staining was performed on P4 and P19 differentiated ALI cultures derived from WT, *Igfbp2*^*−/−*^, and *Ncf1*^*−/−*^ SMGPs. P4 ALI cultures were stained for Bsnd, Foxi1, and Cftr **(Fig. 5A−C)**, whereas P19 ALI cultures were stained for Bsnd, Cftr, and Scgb1a1 **(Fig. 5F−H)**.

**Figure 5.**
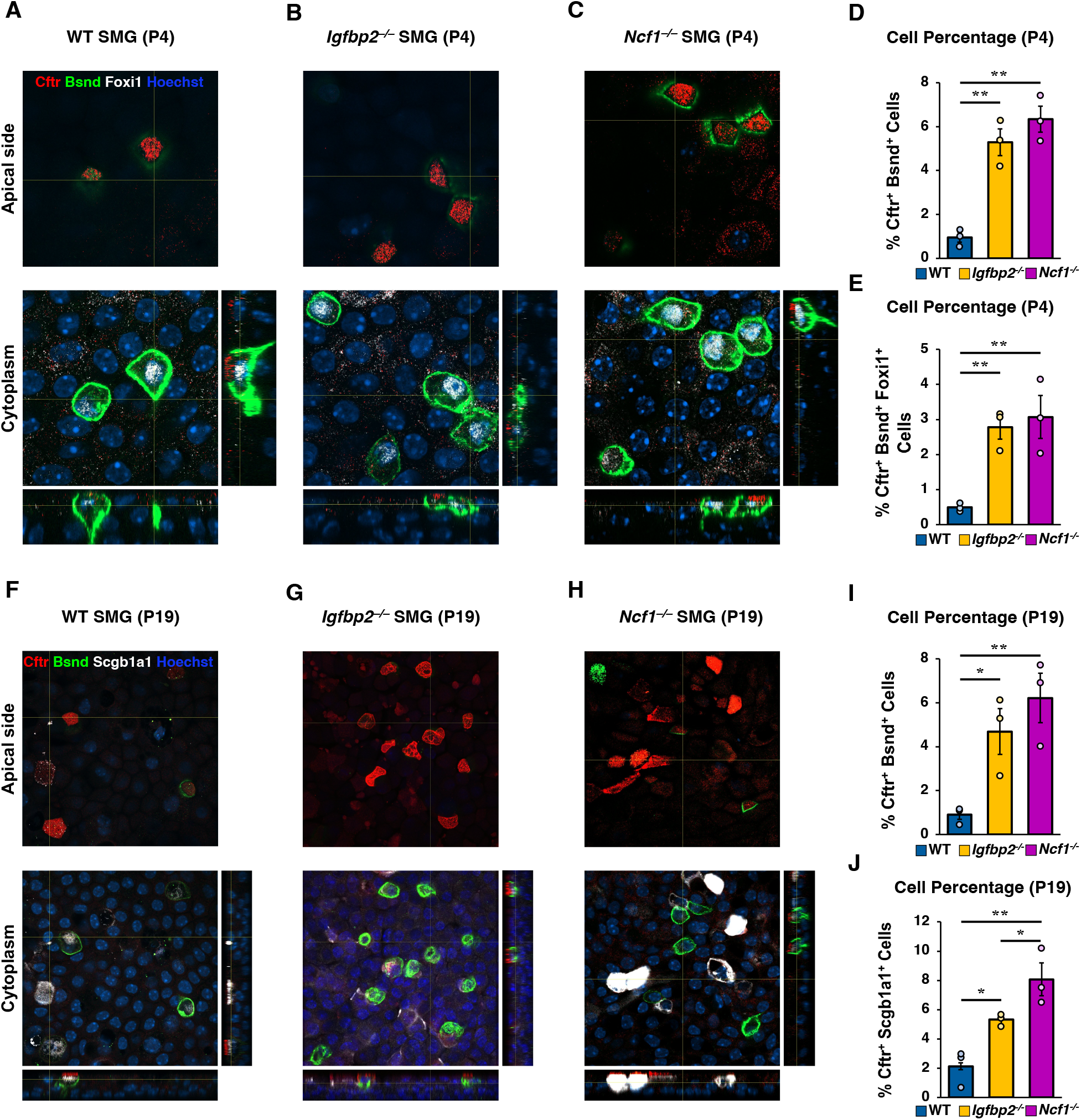
The loss of Ncf1/Igfbp2 signaling positively correlates with the abundance of pulmonary ionocytes, maintained even at high-passage SMGP-derived cultures. Fully differentiated WT, *Igfbp2*^*− /−*^, and *Ncf1*^*−/−*^ SMGP-derived ALI cultures of P4 and P19 were fixed in 4% PFA and underwent the wholemount immunofluorescent staining. **(A-C)** Orthogonal views of 3D images of whole mount ALI membranes of all groups at P4 stained for Cftr (Red), Foxi1 (White), Bsnd (Green), Hoechst (Blue). Quantification of Cftr^+^Bsnd^+^ double positive **(D)**, and Cftr^+^Bsnd^+^Foxi1 triple positive cells **(E)** (n = 3 cultures per genotype, with 1 culture from each of 3 donor groups; each donor group comprised SMGPs from 3 mice). **(F-H)** Orthogonal views of 3D images of whole mount ALI membranes of all groups at P19 stained for Cftr (Red), Scgb1a1 (White), Bsnd (Green), Hoechst (Blue). Quantification of Cftr^+^Bsnd^+^ double positive **(I)**, and Cftr^+^Scgb1a1^+^ double positive cells **(J)** (n=3 donor groups). Data are shown as mean ± SEM. Statistical significance: *P < 0.05, **P < 0.01, ***P < 0.001, ****P < 0.0001; one-way ANOVA with Benjamini−Hochberg FDR post hoc test.

At P4, *Igfbp2*^*−/−*^ and *Ncf1*^*−/−*^ ALI cultures exhibited significantly increased numbers of Cftr^+^Bsnd^+^ double-positive and Cftr^+^Bsnd^+^Foxi1^+^ triple-positive cells relative to WT controls **(Fig. 5D, E)**. Similarly, P19 *Igfbp2*^*−/−*^ and *Ncf1*^*−/−*^ ALI cultures exhibited significantly higher numbers of Cftr^+^Bsnd^+^ and Cftr^+^Scgb1a1^+^ double-positive cells compared to WT cultures **(Fig. 5I, J)**. Collectively, these findings confirm that disruption of Ncf1/Igfbp2 signaling increases pulmonary ionocyte abundance and is also associated with broader alterations in Cftr-expressing secretory cells.

### Loss of Ncf1/Igfbp2 increases ionocyte abundance in tracheal submucosal glands in vivo

To determine whether the increase in ionocyte abundance observed in vitro was recapitulated in vivo, whole-mount IF staining was performed on proximal tracheal tissues from WT, *Igfbp2*^−/−^, and *Ncf1*^−/−^ mice. Tracheae from young (3-month-old) and aged (6-month-old) mice were analyzed to assess the persistence of this phenotype with age. Tracheal submucosal glands were stained for the ionocyte marker Bsnd, and representative orthogonal views of 6-month-old WT, *Igfbp2*^*−/−*^, and *Ncf1*^*−/−*^ tracheae are shown in **Suppl. Fig. S3A−C**. Quantification of Bsnd^+^ cells revealed significantly increased ionocyte abundance in both *Igfbp2*^*−/−*^ and *Ncf1*^*−/−*^ tracheae relative to WT controls at 3 months, which was maintained at 6 months **(Suppl. Fig. S3D)**. These findings demonstrate that loss of Ncf1 or Igfbp2 promotes ionocyte enrichment within tracheal SMGs in vivo, consistent with the phenotype observed in ALI cultures. Collectively, these findings support a role for Ncf1/Igfbp2 signaling in regulating ionocyte specification and/or maintenance and provide in vivo validation of the in vitro observations.

## Discussion

A major challenge in the development of cell-based therapies for cystic fibrosis (CF) is the identification and expansion of airway progenitor populations capable of durable engraftment, epithelial regeneration, and sustained restoration of CFTR function. One key limitation is the progressive decline in progenitor fitness during in vitro expansion, which can compromise epithelial differentiation and reduce CFTR activity [28, 39]. In this study, we identify Ncf1/Igfbp2 signaling as a regulator of airway submucosal gland progenitor fitness, lineage specification, and epithelial function during prolonged expansion. Loss of either Ncf1 or Igfbp2 preserved progenitor characteristics during extended in vitro expansion, enhanced proliferative and organoid-forming capacity, reduced DNA damage accumulation, and promoted expression of DNA repair genes. Disruption of this signaling axis enhanced and preserved Cftr-mediated anion transport in epithelia generated from serially passaged SMGPs, a phenotype associated with enhanced pulmonary ionocyte specification in vitro and increased ionocyte abundance in vivo. Collectively, these findings support a model in which Ncf1/Igfbp2 signaling regulates airway progenitor fitness and epithelial lineage allocation through DNA repair-associated stress pathways rather than telomere erosion.

Loss of Ncf1 or Igfbp2 preserved basal progenitor identity, proliferative capacity, and organoid-forming efficiency during serial passaging, whereas WT cultures showed a progressive decline in these properties **(Fig. 1; Suppl. Figs. S1−2)**. Preservation of progenitor identity in *Ncf1*^−/−^ and *Igfbp2*^−/−^ cultures was accompanied by reduced γ-H2ax accumulation and increased expression of DNA repair genes associated with the Fanconi anemia and homologous recombination pathways, including *Fanca, Fancd2*, and *Rad51*, consistent with enhanced DNA repair capacity during prolonged expansion. These changes were also accompanied by lower cellular ROS at late passage, suggesting that reduced oxidative stress is associated with preservation of progenitor fitness in the *Ncf1*^−/−^ and *Igfbp2*^−/−^ cultures (**Fig. 2**). In contrast, WT cultures exhibited increased *Krt14* and *Pcna* expression at late passage **(Suppl. Fig. S2)** together with higher CellROX fluorescence intensity, consistent with increased cellular stress during prolonged expansion (**Fig. 2**). Given that CellROX Deep Red detects superoxide and hydroxyl radicals (the product of the Fenton reaction between superoxide and Fe^3+^), we hypothesize that disruption of *Ncf1* reduces NADPH oxidase-derived ROS, thereby contributing to the observed functional phenotype. Notably, however, the finding that *Igfbp2*^−/−^ cells also exhibited reduced ROS levels suggests that the relationship between Ncf1, Igfbp2, and cellular oxidative stress may be more complex than a simple linear pathway. A recent study supports a role for Igfbp2 in regulating oxidative stress, as treatment with recombinant Igfbp2 was associated with enhanced oxidative stress and mitochondrial dysfunction in a Parkinson’s disease model [40]. Although this study was performed in a different biological context, these findings are consistent with our observation that *Igfbp2*^−/−^ cells exhibit lower levels of ROS at late passage. Although the underlying mechanism remains unclear, these findings raise the possibility that Igfbp2 may participate in regulating cellular ROS production, redox homeostasis, and DNA repair.

This interpretation is further supported by the stress-associated transcriptional profile of late-passage WT cultures. Prior studies have shown that Krt14 is induced during airway epithelial injury and regeneration and is associated with stress-responsive epithelial remodeling across multiple injury contexts [19, 23, 24]. Similarly, Pcna participates in DNA damage tolerance pathways activated during replication stress [25-27]. The concordance of increased oxidative stress, γ-H2ax accumulation, and *Pcna* expression in late-passage WT cultures further supports a state of progressively increased cellular and replication-associated stress, whereas *Ncf1*^−/−^ and *Igfbp2*^−/−^ cultures maintain progenitor fitness in association with reduced oxidative stress, lower DNA damage accumulation, and enhanced DNA repair capacity.

Previous studies from Reynolds and colleagues demonstrated that repeated airway injury and prolonged stem cell expansion are associated with telomere shortening, depletion of the airway stem cell pool, and reduced regenerative potential [41]. More recently, epithelial stem cells isolated from diseased airways were found to exhibit shortened telomeres and reduced regenerative capacity, further supporting an association between telomere attrition and airway stem cell aging [42]. In contrast, although baseline telomere length differed between genotypes in our study, the rate of telomere attrition during serial passaging was comparable across groups **(Suppl. Fig. S2)**. Instead, preservation of progenitor fitness in *Ncf1*^−/−^ and *Igfbp2*^−/−^ cultures correlated more strongly with reduced DNA damage accumulation and enhanced expression of DNA repair genes. These findings suggest that genome maintenance pathways can preserve long-term airway progenitor function independently of changes in telomere shortening and may represent a distinct mechanism regulating progenitor durability during prolonged expansion.

Preservation of progenitor fitness in *Ncf1*^−/−^ and *Igfbp2*^−/−^ cultures was accompanied by enhanced epithelial function following differentiation, as evidenced by sustained Cftr-mediated chloride transport at both early and late passages relative to WT cultures **(Fig. 3)**. This functional advantage was associated with increased expression of ionocyte markers (*Foxi1, Bsnd*, and *Cftr*) and greater pulmonary ionocyte abundance in vitro and in vivo **(Figs. 4−5; Suppl. Fig. S3)**. Since their identification as a CFTR-enriched epithelial population, pulmonary ionocytes have emerged as important contributors to airway ion transport and epithelial physiology [5, 35-37]. However, efficient generation of ionocytes remains a major challenge in pluripotent stem cell-derived airway differentiation systems [1, 43]. Within this context, our findings identify Ncf1/Igfbp2 signaling as a previously unrecognized regulator of ionocyte specification and suggest that modulation of this pathway may provide a strategy to enhance the generation and maintenance of Cftr-expressing epithelial cell populations. Notably, the effects of Ncf1/Igfbp2 disruption were not restricted to ionocytes, as late-passage knockout cultures exhibited broader alterations in epithelial lineage specification, including enhanced secretory cell differentiation in *Ncf1*^−/−^ cultures and increased ciliated cell lineage marker expression in *Igfbp2*^−/−^ cultures. Together, these findings suggest that Ncf1/Igfbp2 signaling regulates epithelial lineage specification and contributes to the maintenance of Cftr-expressing cell populations during prolonged expansion.

Our findings support a model in which Ncf1/Igfbp2 signaling links genomic stress responses to epithelial lineage specification during progenitor expansion. Disruption of this pathway preserved progenitor fitness, enhanced pulmonary ionocyte abundance, and altered the balance of secretory and ciliated epithelial lineages, resulting in sustained Cftr-mediated ion transport following differentiation. These lineage-specific effects suggest that Ncf1 and Igfbp2 do not simply preserve epithelial differentiation capacity but actively influence epithelial composition. Notably, *Igfbp2*^−/−^ cultures exhibited a relative shift toward a ciliated cell program, whereas *Ncf1*^−/−^ cultures favored secretory differentiation, indicating that distinct components of this signaling axis may differentially regulate epithelial fate decisions. Because ionocytes, secretory cells, and ciliated cells collectively govern airway surface hydration, mucus production, and mucociliary clearance, alterations in epithelial composition may have important consequences for airway homeostasis beyond CFTR function alone. In muco-obstructive diseases such as cystic fibrosis, chronic obstructive pulmonary disease, and chronic bronchitis, therapies that simultaneously enhance airway ion transport, reduce mucus burden, and promote mucociliary clearance could provide substantial therapeutic benefit. Thus, modulation of Ncf1/Igfbp2 signaling may represent a strategy not only to preserve airway progenitor durability during expansion but also to engineer epithelial compositions optimized for functional airway repair.

## Methods

### Mouse models

C57BL/6J mice (The Jackson Laboratory), *Igfbp2*^*−/−*^ mice (gift from Dr. Clifford J. Rosen), and *Ncf1*^*−/−*^ mice (Taconic) were used in this study and previously described [16]. All breeding procedures and experiments were approved by the University of Iowa and the University of Alabama at Birmingham Institutional Animal Care and Use Committee (IACUC).

### CellROX Deep Red ROS Detection Assay

Intracellular ROS was assessed in live WT, *Igfbp2*^−/−^, and *Ncf1*^−/−^ SMGPs at P4 and P19 using CellROX− Deep Red Reagent (Thermo Fisher Scientific, C10422). SMGPs were seeded at 5 × 10^4^ cells/well in chamber slides (Ibidi, 80381). Twenty-four hours after seeding, cells were incubated with CellROX Deep Red for 30 min at 37°C according to the manufacturer’s protocol. Cells were washed with prewarmed PBS and incubated with Hoechst 33342 nuclear stain (Invitrogen) for 10 min in modified SAGM (see supplementary materials). Following Hoechst staining, cells were washed with pre-warmed PBS and maintained in SAGM for live-cell imaging. Fluorescence images were acquired using identical imaging and acquisition settings across genotypes and passages. CellROX Deep Red fluorescence intensity was quantified from acquired images and used as a relative measure of intracellular ROS.

### Short-circuit currents (I_sc_) measurements

Transepithelial short-circuit current (*I*_*sc*_) was measured using a self-contained Ussing chamber system (Physiologic Instruments) as previously described [44]. For chloride conductance, epithelia were bathed with 37°C symmetrical chloride buffer containing 135 mM NaCl, 2.4 mM K_2_HPO_4_, 0.6 mM KH_2_PO_4_, 1.2 mM CaCl_2_, 1.2 mM MgCl_2_, 10 mM dextrose, and 5 mM Hepes (pH 7.4) and bubbled with air. *I*_*sc*_ was recorded following sequential apical addition of amiloride (100 µM), DIDS (100 µM), IBMX (100 µM) plus Forskolin (10 µM), and GlyH101 (50 µM) plus CFTR_Inh_-172 (50 µM).

### Statistical analysis

Statistical analyses were performed using GraphPad Prism 7 software and Excel, Electrophysiological line graphs were assembled using Excel. Exact P values and statistical test performed are provided in the figure legends. Error bars indicate the standard error of the mean (SEM).

## See Supplemental Materials for details of the following areas

Animals; Tissue processing and cell harvesting; Air-liquid interface cultures; Organoid cultures; Reverse transcription and quantitative polymerase chain reaction; Preparation of total cell lysates and Western blot analysis; Absolute telomere length quantification; EdU incorporation assay; Immunofluorescent staining of whole mount ALI transwell membranes; Immunofluorescent staining of whole mount mouse tracheae; Imaging and image processing.

## Author contributions

JH: conception, experimental design, acquisition, analysis, interpretation, writing, revision; KDC: acquisition, analysis, interpretation, revision; OA: acquisition, analysis, interpretation; GF: acquisition, analysis; CAY: analysis, interpretation, revision; WSS: conception, experimental design, acquisition, analysis, interpretation, revision, editing, supervision; JFE: conception, interpretation, revision, editing, funding acquisition, supervision.

## Data availability statement

The data generated in this study are available in the published article and its supplementary files; additional data are available from the corresponding authors upon reasonable request.

