## Supplementary Materials for "Ncf1/Igfbp2 Signaling Regulates Airway Submucosal Gland Progenitor Fitness and Ionocyte Specification"

**Supplemental files**

Supplemental Materials and Methods

Supplemental Figures

Supplemental Table

### Supplemental Materials and Methods

### Materials and Methods

*Animals*

Two-, three-, and six-month-old C57BL/6J mice (The Jackson Laboratory), *Igfbp2*$^{-/-}$ mice (a generous gift from Dr. Clifford J. Rosen), and *Ncf1*$^{-/-}$ mice (Taconic) were used in this study and previously described [1]. Animals were group housed and maintained in a 12-h light/dark cycle with set points at 73°F and 50% relative humidity. Animals were grouped according to genotype without randomization. Male and female mice were equally represented within each genotype group; sex was not considered an experimental variable. All breeding procedures and experiments were performed according to a protocol approved by the University of Iowa and the University of Alabama at Birmingham Institutional Animal Care and Use Committee (IACUC).

*Tissue processing and cell harvesting*

SMGPs were isolated from resected tracheae of two-month-old mice using a sequential enzymatic digestion protocol as previously described [2]. Tracheae were opened longitudinally to expose the surface airway epithelium (SAE) and incubated in 1.5 mg/mL pronase (Roche) in DMEM/F12 at 37°C for 1 hour with gentle agitation. Following digestion, tissues were washed five times with DMEM/F12, and the SAE was removed by repeated gentle pipetting using a 5 mL plastic pipette. The remaining tracheal tissue was processed for SMG cell isolation. Tissues were minced into fragments $\leq 3$ mm³ using fine surgical scissors and incubated in 2× collagenase/hyaluronidase solution (StemCell Technologies) diluted in DMEM/F12 at 37°C for 45 minutes with gentle nutation. Pre-warmed 0.25% trypsin-EDTA (Life Technologies) was then added to achieve a final concentration of 0.05% and incubated for an additional 30 minutes at 37°C with gentle agitation. The cell suspension was mechanically dissociated by pipetting and filtered through a 100 μm cell strainer to obtain a single-cell suspension. Cells were collected by centrifugation and resuspended in DMEM/F12, followed by a medium exchange into modified SAGM™ (Small Airway Epithelial Cell Growth Medium BulletKit, Lonza), consisting of SABM™ basal medium (Cat# CC-3119) supplemented with SAGM™ SingleQuots® additives (Cat# CC-4124) with modifications as previously described [3]. The medium was further supplemented with small-molecule modulators, including the BMP pathway inhibitor DMH-1, the WNT agonist CHIR99021, and the TGF-β receptor inhibitor A83-01 (each at a final concentration of 1 μM), as well as the ROCK inhibitor Y-27632 (2 μM), all prepared from 10 mM stock solutions. Penicillin–streptomycin was added at 1% (5 mL per 500 mL of complete medium). Cells were washed once in modified SAGM™ and subsequently plated in the same medium for culture. All centrifugation steps were performed at 250 × g for 5 minutes.

*Air-liquid interface (ALI) cultures*

0.33 $cm^2$ polyester transwell membranes (Corning) were pre-treated with 804G-conditioned media, after which the expanded primary WT, *Igfbp2*$^{-/-}$, and *Ncf1*$^{-/-}$ SMG progenitors were seeded in modified SAGM (described above) at a density of $2 \times 10^5$ cell. 24 hours post-seeding, the modified SAGM media was replaced with Pneumacult ALI medium (StemCell Technologies, Inc.) that was added to the basolateral side of the transwells in order to establish the air-liquid interface. ALI cultures were maintained for 7 weeks to permit complete differentiation of SMGPs prior to their use in downstream experiments.

*Organoid cultures*

Organoid cultures were established to assess OFE. Wells of a 48-well glass-bottom plate (MATTEK, P48G-1.5-6-F) were coated with 15 µL of Matrigel (Corning) and incubated at 37°C for 30 minutes prior to cell seeding. WT, *Igfbp2*$^{-/-}$, and *Ncf1*$^{-/-}$ SMGPs at P4 and P19 were seeded at a density of $2 \times 10^3$ cells per well in 35 µL of modified SAGM medium (described above) mixed 1:1 with Matrigel. Plates were further incubated at 37°C for 15 minutes to allow Matrigel polymerization, after which pre-warmed modified SAGM was added. Organoid cultures were maintained in modified SAGM for 9 days prior to imaging and OFE quantification. For live-cell staining, organoid cultures were incubated for 30 minutes prior to imaging with Calcein AM (Biotium, #80011-3) prepared from a 1 mg/mL stock solution in DMSO and diluted 1:500 in culture medium. Imaging was performed using the live-cell imaging chamber of a Nikon confocal microscope. The entire well area containing organoids was captured for each culture, and organoid counts were quantified using NIS-Elements software. OFE was calculated as (organoid count / seeding density) × 100.

*Reverse transcription and quantitative polymerase chain reaction (qRT-PCR)*

Seven weeks post-differentiation in Pneumacult ALI medium, ALI cultures were harvested for RNA extraction. cDNA was synthesized by reverse transcription and subsequently analyzed by qRT-PCR. RNeasy Plus Mini Kit (Qiagen) was used to extract total RNA and was converted into cDNA using the SuperScript IV VILO Master Mix Kit (ThermoFisher Scientific) according to the protocol provided by the manufacturer. TaqMan Universal Master Mix II with UNG (Applied Biosystems) was used to quantify the transcription levels of our target genes relative to β-Actin (internal control) according to the manufacturer's protocol. We used the following gene expression assays for qPCR analysis, showed in Figure 6: Mm02619580_g1 Actb 4448484 TaqMan Gene Expression Assay, SM VIC PL for β-Actin, Mm00458451_m1 Foxi1 4331182 TaqMan Gene Expression Assay, SM for Foxi1, Mm00445197_m1

Cftr 4331182 TaqMan Gene Expression Assay, SM for Cftr and Mm00475532_m1 Bsnd 4331182 TaqMan Gene Expression Assay, SM for Bsnd, Mm01267279_m1 FoxJ1 4331182 TaqMan Gene Expression Assay, SM for FoxJ1, and Mm00600221_m1 Spdef 4331182 TaqMan Gene Expression Assay, SM for Spdef (ThermoFisher Scientific). The sequences of the primers and probes for qPCR analysis in Figures 1 and 3 are provided in **Suppl. Table S1**.

*Preparation of total cell lysates and Western blot analysis*

WT, *Igfbp2*$^{-/-}$, and *Ncf1*$^{-/-}$ SMGPs at passages 4 (P4) and 19 (P19) were plated at a density of 1 × $10^6$ cells per 100 mm tissue culture dish and maintained for 5 days in modified SAGM™ (see above). Following culture, cells were collected into 15 mL conical tubes and pelleted by centrifugation. Cell pellets were lysed in RIPA buffer (Sigma) supplemented with phosphatase inhibitor mini tablets (Pierce) and cOmplete protease inhibitor cocktail (Roche) and incubated on ice for 20 minutes. Lysates were sonicated for 20 seconds and centrifuged at 13,000 × g for 5 minutes at 4°C to remove debris. The resulting supernatant was transferred to screw-cap tubes, snap-frozen in liquid nitrogen, and stored at −80°C. Protein concentration was determined using the Bio-Rad protein assay according to the manufacturer's instructions. Equal amounts of protein (150 µg per sample) were denatured in Laemmli sample buffer (Sigma) by boiling and resolved on 4–20% gradient SDS-PAGE gels. Proteins were transferred to nitrocellulose membranes, which were subsequently blocked in 2% bovine serum albumin (BSA) prepared in filtered 1× TBS buffer (36 mM Tris base, 50 mM NaCl, 0.5% Tween 20). Membranes were incubated overnight at 4°C with primary antibodies diluted in 0.5% BSA in 1×TBS, including rabbit monoclonal anti-γ-H2ax (Abcam, 1:1000), goat polyclonal anti-Gapdh (Thermo Fisher, 1:1000), and chicken polyclonal anti-cytokeratin 5 (BioLegend, cat#905901, 1:1000). After washing, membranes were incubated for 2 hours at room temperature with secondary antibodies diluted in 0.5% BSA in 1×TBS: IRDye 680 donkey anti-rabbit, IRDye 800 donkey anti-goat antibodies (LI-COR Biosciences, 1:10,000), and Alexa Fluor 488–conjugated donkey anti-chicken (Jackson ImmunoResearch, 1:10,000). Fluorescent signals were detected using an Odyssey M imaging system (LI-COR Biosciences), and band intensities were quantified using Empiria Studio software (LI-COR Biosciences).

*Absolute telomere length quantification*

Absolute telomere length was quantified using the Absolute Mouse Telomere Length Quantification qPCR Assay Kit (ScienCell, M89180). WT, *Igfbp2*$^{-/-}$, and *Ncf1*$^{-/-}$ SMG progenitors at passages 4, 10, 15, and 19 were detached using Accutase (STEMCELL Technologies), followed by genomic DNA extraction using the DNeasy Blood & Tissue Kit (QIAGEN). Quantitative PCR and telomere length quantification were subsequently performed according to the manufacturer's instructions.

*EdU incorporation assay*

Cell proliferation was assessed using the Click-iT™ EdU Cell Proliferation Kit for Imaging, Alexa Fluor™ 488 dye (Thermo Fisher Scientific, C10337). WT, *Igfbp2*$^{-/-}$, and *Ncf1*$^{-/-}$ SMG progenitors at passages 4 and 19 were seeded at a density of $5 \times 10^4$ cells per well in 3-well chamber slides (Ibidi, 80381). Twenty-four hours after seeding, cells were pulsed with 10 μM EdU prepared according to the manufacturer's instructions for 3 hours, followed by three washes with modified SAGM medium (described above).

*Immunofluorescent staining of whole mount ALI transwell membranes*

Whole mount ALI cultures were fixed in 4% PFA for 1 hour, followed by three 5-minute washes in PBS. Washed membranes were boiled at 98˚C for 35 minutes in an antigen retrieving citrate buffer containing 10 mM sodium citrate, 0.05% Tween 20, and adjusted to pH 6.0. After gradual cooling down to room temperature, membranes were washed by three 5-minute PBS washes. Membranes were then incubated in blocking buffer containing 20% normal donkey serum, 0.3% Triton X-100, and 1 mM $CaCl_2$ in PBS for 2 hours at room temperature. Membranes were incubated overnight at room temperature with a cocktail of primary antibodies targeting the proteins of interest, including rabbit anti-BSND (1:300; Abcam, Cat# ab196017), mouse anti-CFTR (1:150; Abcam, Cat# ab2784), goat anti-SCGB1A1 (1:4000; R&D Systems, Cat# ABS1673), and goat anti-FOXI1 (1:100; Abcam, Cat# ab20454). Antibodies were diluted in PBS-based diluent buffer containing 1% normal donkey serum, 0.3% Triton X-100, and 1 mM $CaCl_2$. Following incubation, membranes were washed three times in PBS for 5 minutes each and then incubated for 2 hours at room temperature with fluorescently conjugated secondary antibodies diluted 1:250 in diluent buffer containing 1% normal donkey serum, 0.3% Triton X-100, and 1 mM $CaCl_2$ in PBS. Secondary antibodies included Alexa Fluor 488 donkey anti-rabbit IgG (Jackson ImmunoResearch, Cat# 711-547-003), Alexa Fluor 647-conjugated donkey anti-goat $F(ab')_2$ fragment IgG (Jackson ImmunoResearch, Cat# 705-606-147), and Alexa Fluor 594-conjugated donkey anti-mouse $F(ab')_2$ fragment IgG (Jackson ImmunoResearch, Cat# 715-586-151). Nuclear counterstaining was concurrently performed using Hoechst 33342 (Invitrogen) diluted in the same buffer. Membranes were washed by three 5-minute PBS washes, followed by mounting on a slide with Aqua-Mount (Thermo Scientific) and flattening between two magnets for at least an hour before imaging.

*Immunofluorescent staining of whole mount mouse tracheae*

Tracheae from three- and six-month-old WT, *Igfbp2*$^{-/-}$, and *Ncf1*$^{-/-}$ mouse donors were harvested, cut open longitudinally, and fixed in 4% PFA for 2 hours, followed by three 30-minute PBS washes at room temperature on a rocker. Tracheae were heated in an antigen retrieving citrate buffer containing 10 mM sodium citrate, 0.05% Tween 20, and pH 6.0, overnight at 55˚C with gentle rocking. Tracheae were then washed twice for 30 minutes with pre-warmed PBS with rocking, followed by an overnight incubation at 37˚C with gentle rocking in blocking buffer containing 20% normal donkey serum, 0.3% Triton X-100, and 1 mM $CaCl_2$ in PBS. Tracheae were incubated for 2 days at 37˚C with a rabbit anti-BSND (1:300; Abcam, Cat# ab196017) primary antibody in diluent buffer containing 1% normal donkey serum, 0.3% Triton X-100, and 1 mM $CaCl_2$ in PBS. Tracheae were washed twice for 30 minutes with pre-warmed PBS with rocking, followed by incubation with Alexa Fluor 488 donkey anti-rabbit IgG (Jackson ImmunoResearch, Cat# 711-547-003) secondary antibody and Hoechst 33342 (Invitrogen) in diluent buffer overnight at 37°C with rocking. Tracheae were mounted on a slide with Aqua-Mount (Thermo Scientific) and flattened between two magnets.

*Imaging and image processing*

Immunofluorescence (IF) images presented in Figures S1 and 2 were acquired using a Nikon AX confocal microscope using a ×20 objective, with 2× digital zoom for CellROX images and no zoom for EdU images. Image acquisition was controlled using Nikon NIS-Elements AR software. IF images presented in Figures 5 and S3 were acquired using a line-scanning confocal microscope (Zeiss LSM-980, Carl Zeiss, Germany) using ×20 and ×63 objectives and controlled using ZEN 2010 acquisition software. Images were processed and quantified using the offline MetaMorph software (Molecular Devices) and ImageJ. Quantification of the numbers of cells positive and negative for specific markers was achieved by using the multi-wavelength cell scoring application module of the MetaMorph software. The following parameters were set on the multi-wavelength cell scoring application module in order to quantify cells positive or double-positive for our target markers: the average nucleus and cytosol sizes; whether a marker is expressed in the nucleus, cytosol, or both; and the intensity above local background and the percentage of the cell that must be stained for it to be considered positive for a certain marker.

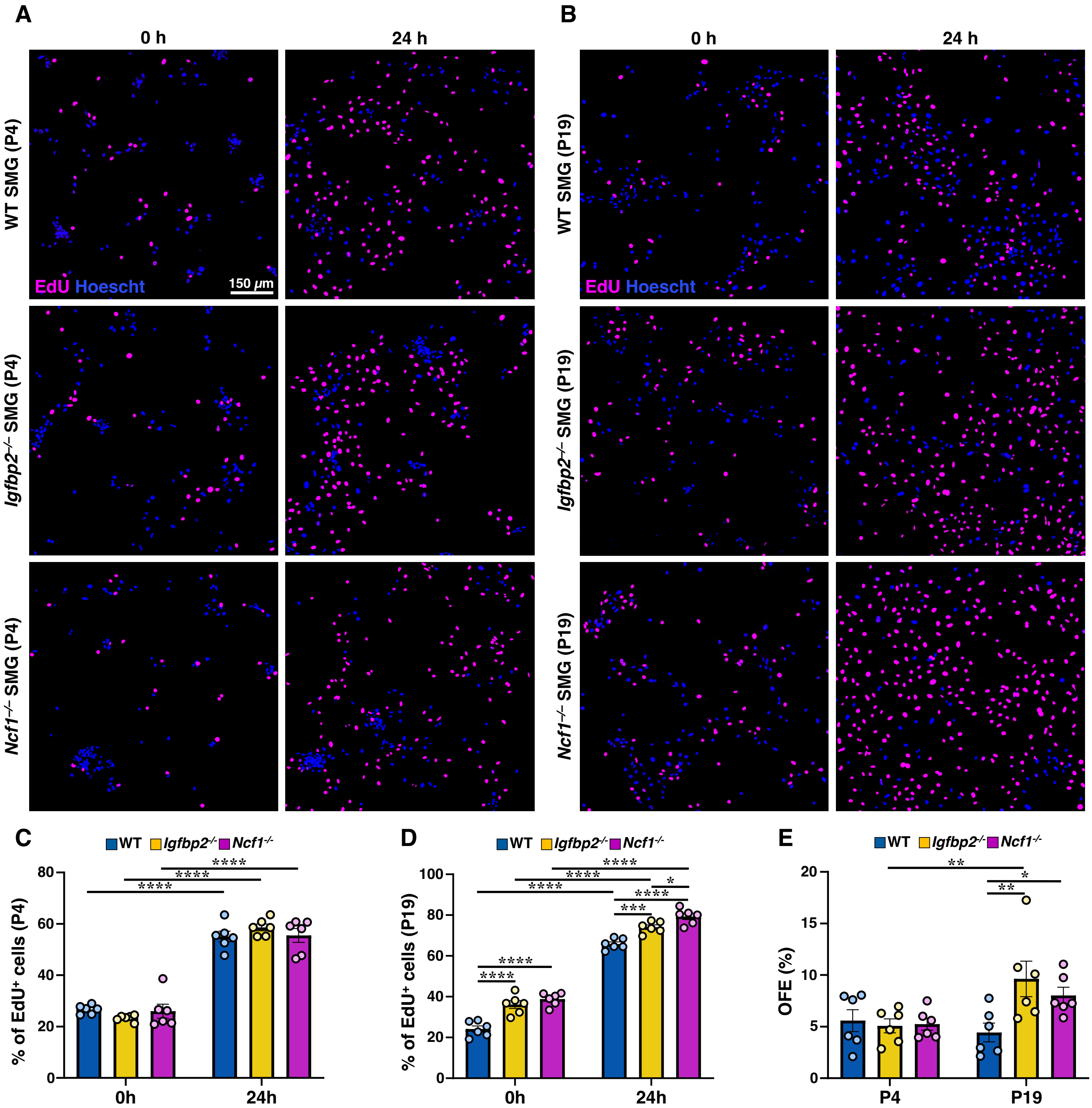


**Supplementary Figure S1. *Ncf1*$^{-/-}$ and *Igfbp2*$^{-/-}$ SMG progenitors retain proliferative capacity and organoid-forming potential during long-term culture. (A, B)** Representative images of EdU incorporation in WT, *Igfbp2*$^{-/-}$, and *Ncf1*$^{-/-}$ SMG progenitors at passage 4 (A) and 19 (B) following a 3-hour pulse and 0h or 24h chase. **(C, D)** Quantification of the percentage of EdU-positive cells at P4 (C) and P19 (D) (n = 6 cultures per genotype, with 2 cultures from each of 3 donor groups; each donor group comprised SMGPs from 3 mice). **(E)** Organoid-forming efficiency (OFE) of WT, *Igfbp2*$^{-/-}$, and *Ncf1*$^{-/-}$ SMG progenitors at P4 and P19 (n = 6 cultures per cell type). Data are shown as mean ± SEM. Statistical significance: *P < 0.05, **P < 0.01, ***P < 0.001; one-way ANOVA with Benjamini–Hochberg FDR post hoc test.

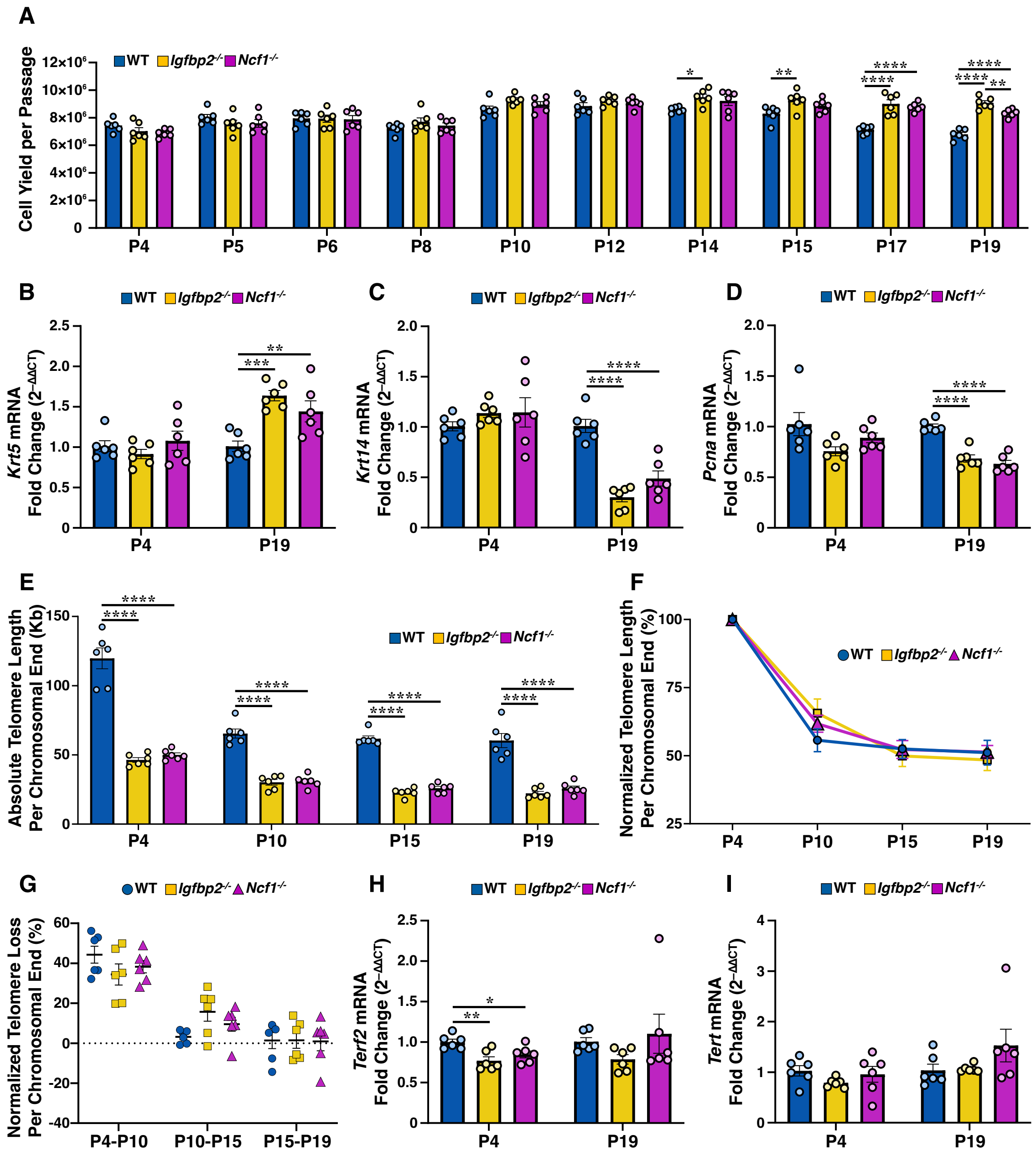


**Supplementary Figure S2. *Ncf1*$^{-/-}$ and *Igfbp2*$^{-/-}$ SMG progenitors sustain cell expansion and basal marker expression with altered stress-associated gene profiles and unchanged telomere attrition rate. (A)** Cell yield per passage from P4 to P19 in WT, *Igfbp2*$^{-/-}$, and *Ncf1*$^{-/-}$ SMG progenitors (n = 6 cultures per cell type). **(B–D)** qPCR analysis of *Krt5* (B), *Krt14* (C), and *Pcna* (D) expression at P4 and P19 (n = 6 cultures per cell type). Gene expression was normalized to *Gapdh*, and relative expression was calculated using the $2^{-\Delta\Delta CT}$ method, with ΔΔCT values determined relative to the mean ΔCT of WT controls. **(E)** Absolute telomere length per chromosomal end (kb) at P4, P10, P15, and P19. **(F)** Telomere length normalized to P4 baseline (%). **(G)** Quantification of normalized telomere length loss per chromosomal end across passages (P4–P10, P10–P15, P15–P19). **(H, I)** qPCR analysis of *Terf2* (H) and *Tert* (I) expression at P4 and P19 (n = 6 cultures per genotype, with 2 cultures from each of 3 donor groups; each donor group comprised SMGPs from 3 mice). Data are shown as mean ± SEM. Statistical significance: *P < 0.05, **P < 0.01, ***P < 0.001, ****P < 0.0001; one-way ANOVA with Benjamini–Hochberg FDR post hoc test.

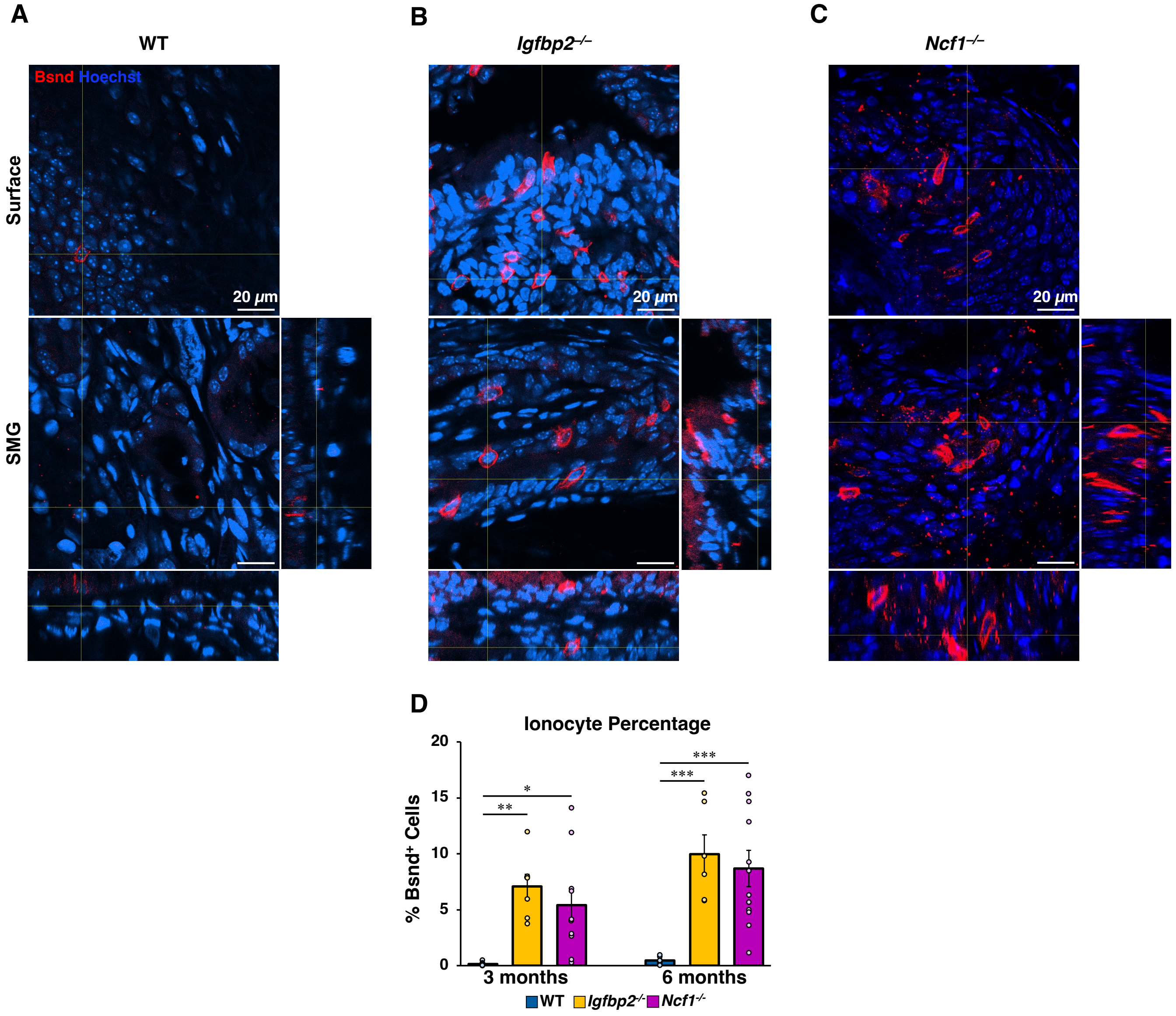


**Supplementary Figure S3. Disruption of Ncf1/Igfbp2 signaling promotes and maintains ionocyte specification in airway submucosal glands.** WT, *Igfbp2*$^{-/-}$, and *Ncf1*$^{-/-}$ mice were euthanized at three and six months of age. Tracheae were harvested and processed for whole-mount immunofluorescence staining for Bsnd (green) and Hoechst (blue), followed by confocal imaging of submucosal glands. **(A–C)** Representative images show glandular localization of the ionocyte marker Bsnd$^{+}$ cells within the airway epithelium of six-month-old mouse tracheas. **(D)** Quantification of Bsnd$^{+}$ ionocytes in three- and six-month-old mice (WT: n = 6 mice at 3 months and n = 6 mice at 6 months; *Igfbp2*$^{-/-}$: n = 7 mice at 3 months and n = 6 mice at 6 months; *Ncf1*$^{-/-}$: n = 10 mice at 3 months and n = 12 mice at 6 months). Data are shown as mean ± SEM. Statistical significance: *P < 0.05, **P < 0.01, ***P < 0.001, ****P < 0.0001; two-way ANOVA with Benjamini–Hochberg FDR post hoc test.

### Supplemental Table

**Supplementary Table S1.** Sequences of primers and probes for quantitative PCR.

| Gene Name | **Forward Primer Sequence** | **Reverse Primer Sequence** | **Probe Sequence** |
|---|---|---|---|
| *Rad51* | 5'-GGC AGC GAT GTC CTA GAT AAT G-3' | 5'- CAG TGC ATA CCT GGA TTC TAC C-3' | 5'-/56-FAM/TCC TTT ACC/ZEN/AAG CGT CAG CCA TGA /31ABkFQ/-3' |
| *Fancd2* | 5'-TCT GAA GGC CAT AGA GGA GAT-3' | 5'- CTG GTC AAC GTA GGG AAT GTA G-3' | 5'-/56-FAM/ACT GGT GTT/ZEN/GGT GTC CCA GAA CTG/31ABkFQ/-3' |
| *Fanca* | 5'- CTT TGC GTG GAG AGG AGT TAG-3' | 5'-AAA GAA GAA GAG GGC GAT GAG-3' | 5'-/56-FAM/AAA GGC TGG/ZEN/ACT GCC AAA GAG AGG/31ABkFQ/-3' |
| *Terf2* | 5'-GAA GAG GAC CAG CTG TTT GA -3' | 5'-ACC CAC TCG CTT TCT TCT ATG-3' | 5'-/56-FAM/AGT TCA GGC/ZEN/ACC AGG TGA AGA CAG /3IABkFQ/-3' |
| *Tert* | 5'-GAG AGA GAG AGT CAG TCA GTC A-3' | 5'-TCA GAG AAG GCA TGG AAC ATC-3' | 5'-/56-FAM/TGA TGA AGT/ZEN/CCA CAG GGA TCA CGC/3IABkFQ/-3' |
| *Pcna* | 5'- CAC ATT GGA GAT GCT GTT GTG-3' | 5'-GTT ACC GCC TCC TCT TCT TTA T-3' | 5'-/56-FAM/AAG TGG AGA /ZEN/GCT TGG CAA TGG GAA /3IABkFQ/-3' |
| *Krt14* | 5'-GAG CGG CAA GAG TGA GAT TT-3' | 5'- CTT TGG TCT CCT CCA GGT TAT TC-3' | 5'-/56-FAM/AGT CCC AGC/ZEN/TCA GCA TGA AAG CAT/31ABkFQ/-3' |
| *Krt5* | 5'- CAA CAA GTT TGC CTC CTT CAT C-3' | 5'- CGG ATC CAG GTT CTG CTT TAT -3' | 5'-/56-FAM/TTC CTG GAG/ZEN/CAG CAG AAC AAG GTC/31ABkFQ/-3' |
| *Gapdh* | 5'-GGA GAA ACC TGC CAA GTA TGA -3' | 5'-TCC TCA GTG TAG CCC AAG A -3' | 5'-/56-FAM/TCA AGA AGG/ZEN/TGG TGA AGC AGG CAT/31ABkFQ/-3' |
